# Reannotation of the zebra finch genome reveals undiscovered transcriptional complexity

**DOI:** 10.1101/2024.01.09.574630

**Authors:** Hugues Richard, John Wiedenhoeft, Constance Scharff, Iris Adam

## Abstract

The zebra finch has become a paradigmatic model system to mechanistically investigate vocal communication from genes to behavior. Here, we report 1) a comprehensive map of the polyadenylated fraction of the transcriptome of the male zebra finch telencephalon at two ages, during the song learning phase at 50 days and when fully adult at 2 years as well as 2) as an expanded and refined annotation of the zebra finch genome based on our transcriptome data. Using high‐throughput next generation sequencing of paired‐end strand‐specific cDNA fragments, we detected over 50% of the annotated protein coding transcripts (ENSEMBL v.55) in the telencephalon at both stages. We exploited the information gained from the paired‐end sequence reads and from reads falling onto splice junctions to update the existing annotation (ENS55) with new transcript structures and alternative splicing events. The dataset allowed to refine 2822 of the available gene models by extending them in the 3’‐UTR and to detect 11391 undiscovered transcriptional units in non‐coding regions. Our results illustrate the value of continuously incorporating new transcriptional evidence into existing annotations.

## INTRODUCTION

During the last decades, the Australian zebra finch (*Taeniopygia castanotis*) has become the best suited model system to study the neural and peripheral substrates of vocal learning and production, brain sexual dimorphism and adult neurogenesis. Recent advances, like the sequencing of the genome[1, 2], virally mediated manipulation of gene expression [3–7] and transgenic zebra finches [8, 9] have expanded the experimental possibilities into the genetic realm.

In the past decade, high throughput RNA‐sequencing (RNA‐seq) using next generations sequencing methods (NGS) has yielded unprecedented insight into the molecular mechanisms of life from prokaryotes to humans including many non‐model species. The main advantage of RNA‐seq, next to a large dynamic range, is that expression of transcripts can be discovered and quantified without a priori knowledge. However, the most prevalent workflow is to align sequenced reads to a reference genome and quantify transcript expression by quantifying the number of reads residing within the gene annotation of the reference genome. Depending on the quality of the annotation, this step can introduce strong biases due to incomplete annotation. Because gene annotation is often relies on detection of protein coding regions, information on noncoding regions—like the 5’‐ and 3’‐untranslated regions (UTR)—and alternative splice forms can be very incomplete.

To advance the use of the zebra finch as genetic model organism we previously generated transcriptome data to test the completeness, expand and refine the Ensembl annotation version 55 (ENS55). We discovered that the majority of ENS55 gene models were lacking annotation of the 3’UTR as well as alternative splice forms. We developed and employed a reannotation strategy using our experimental data to precisely map polyadenylation sites, extend existing and add new gene models to the existing annotation. Furthermore, we performed PCR validation of the improved annotation on the original RNA as well as new independent samples. Overall, our results illustrate the value of incorporating new experimental data into exiting annotations.

## MATERIAL AND METHODS

### Animal husbandry

All zebra finches originated from the captive breeding colony at Freie Universität Berlin (Permit Number: ZH147, ZH144). Adult individuals were housed in free flight aviaries under a 12:12h light:dark‐cycle with food and water provided *ad libitum*. Breeding pairs were kept in cages. Chicks were removed from their home cages 90 days post hatching (PHD90).

### Organ dissection and storage

After decapitation, brains were quickly dissected and the forebrain was separated from the rest of the brain. Tissue was snap frozen in liquid nitrogen after dissection and stored at −80 °C.

### RNA extraction

Forebrain tissue was either pooled by age (transcriptome sequencing) or treated individually (PCR validation). Total RNA was extracted from each sample using the Trizol (Invitrogen) following the manufacturer’s protocol. Samples used for PCR confirmation were treated with DNAse (Turbo DNA free, Ambion) following the manufacturer’s recommendations.

### cDNA‐synthesis

For PCR validation of the re‐annotation, total RNA was reverse transcribed using 1 µg of RNA, 200 ng random hexamer primers (QPCR) or 200 ng oligo‐dT primers, 10 nmol of each dNTP, 50 mM DTT, 200 U SuperScript III (Invitrogen) and 40 U RNasinPlus (Promega). The temperature program was chosen as follows: 5’ at 65 °C, cool down on ice, 5’ at 25 °C (if random hexamer primers were used), 45’ at 50 °C, 15’ at 72 °C. Reactions without reverse transcriptase (‐RT) were always included to control for genomic DNA contaminations.

### Polymerase chain reaction (PCR)

Non‐quantitative PCR reactions were carried out in a reaction volume of 25 µL containing 0.18 mM dNTPs, 2 mM MgCl_2_, 70 mM TrisHCl, 17.25 mM ammonium sulfate, 0.1 % Tween20, 0.32 nM forward primer, 0 32 nM reverse primer and 2 U Taq‐polymerase. All primer pairs and their annealing temperature used in non‐quantitative PCR assays are listed in **Table 1**. No template controls (NTC) to detect DNA contaminations were always included. The PCR was typically run using the following temperature program: 3’ at 94 °C, 30 cycles each with 30’’ at 94 °C, 30’’ at annealing temperature (see Table 1), 45’’ at 72 °C followed by 5’ at 72 °C. The elongation time was extended to up to 5’ for long amplicons. If PCR products were intended for cloning into plasmids, a Taq polymerase with proof reading activity was used (e.g. Phusion High Fidelity DNA polymerase, NEB). PCR products were separated on a 1–3 % TBE agarose gel.

**Table 1:**
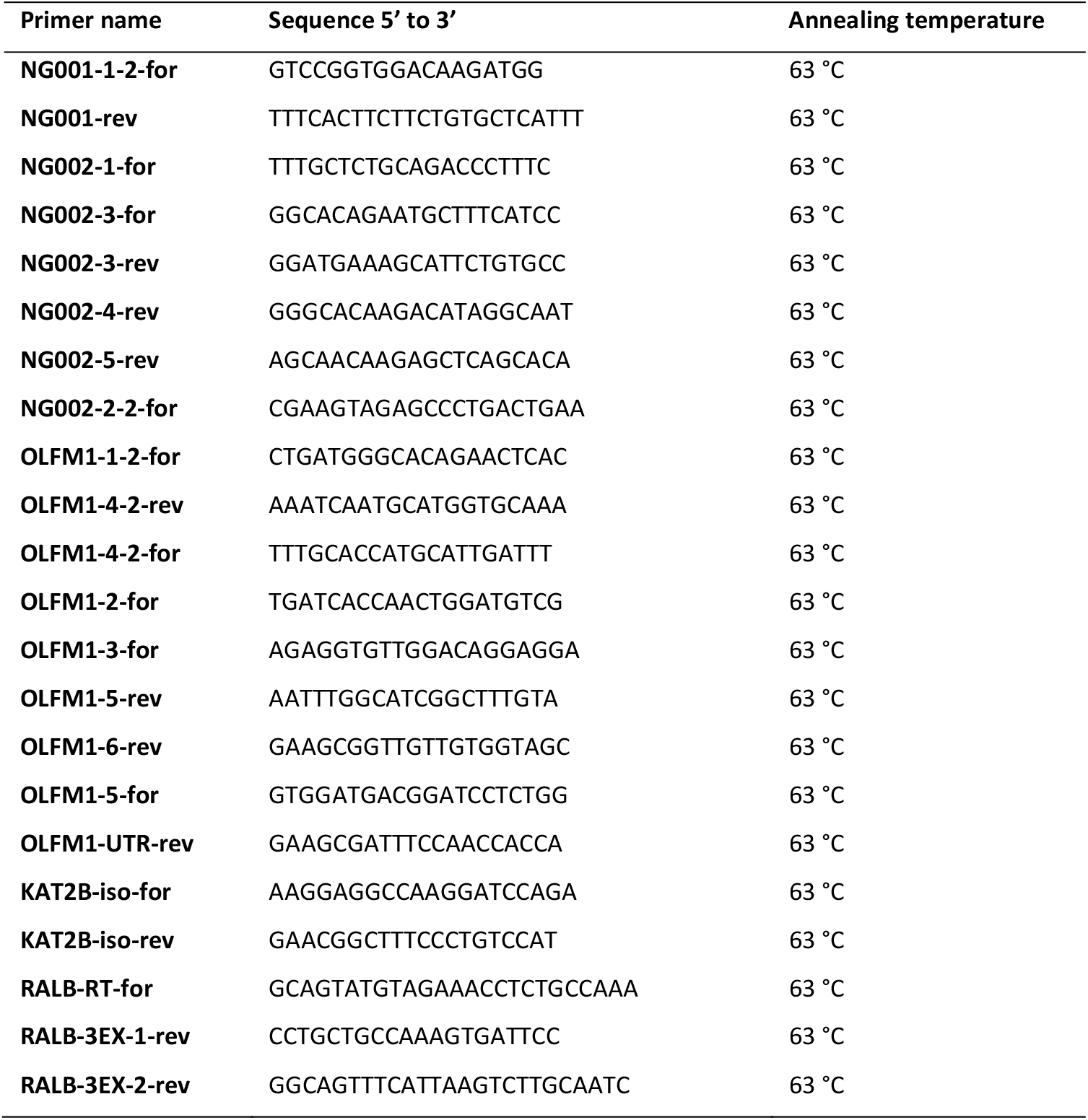
List of primers used for non‐quantitative PCR.

### Transcriptome sequencing

Generation of the RNA‐seq data was previously described [1] and is briefly repeated here.

### Library preparation and sequencing

#### PolyA^+^RNA purification

PolyA^+^RNA was purified with the Dynabeads mRNA purification kit (Invitrogen) following the manufacturer’s instructions, and treated with 0.2 U of TURBODNase (Ambion) per 1 μg of RNA for 30’ at 37 °C.

#### First strand synthesis (FSS)

FSS reactions was prepared by mixing 0.5 μg of polyA^+^RNA, 40 ng of random hexamer primers (Invitrogen) and 25 pmol of oligo‐dT primer (Invitrogen) in 8.5 μL of 1x reverse transcription buffer (Invitrogen), 0.5 mM dNTPs, 5 mM MgCl_2_ and 10 mM DTT. The mixture was incubated at 98 °C for 1’, then at 70 °C for 5’ and subsequently cooled to 15 °C at 0.1 °C/s. At 15 °C, 0.5 μl actinomycin D solution (120 ng/μL), 0.5 μL RNaseOUT (40 U/μL, Invitrogen) and 0.5 μL SuperScript III polymerase (200 U/μL, Invitrogen) were added to the reaction. The temperature of the reverse transcription reaction was increased heating from 15 to 25 °C at 0.1 °C/s, incubation at 25 °C for 10’; heating from 25 to 42 °C at 0.1 °C/s, incubation at 42 °C for 45’; heating from 42 to 50 °C at 0.1 °C/s; incubation at 50 °C for 25’. Reverse transcriptase activity was finally inactivated at 75 °C for 15’.

#### Removal of dNTPs

20 μL EB (10 mM TrisCl, pH 8.5, Qiagen) were added to the reaction. dNTPs were removed by purification of the first strand mixture on a self‐made 200 μL G‐50 gel filtration spin‐column equilibrated with 1 mM TrisHCl, pH 7.0.

#### Second strand synthesis (SSS)

Water was added to the purified FSS reaction to bring the final volume to 55.5 μL. The mixture was cooled on ice. Then, 19.5 μL the “second strand mixture” (1.5 μL 10 mM dNTP mix; 15 μL 5x SSS buffer (Invitrogen); 0.5 μL of *E. coli* ligase (10 U/μL, NEB); 2 μL of DNA polymerase I (10 U/μL, NEB) and 0.5 μL RNase H (2 U/μL, Invitrogen)) were added. SSS reactions were incubated at 16 °C for 2 h. Double‐stranded cDNA was purified on QIAquick columns (Qiagen) according to the manufacturer’s instructions.

#### DNA fragmentation

Roughly 250 ng of double‐stranded cDNA were fragmented by sonication with a UTR200 (Hielscher Ultrasonics GmbH) using the following conditions: 1 h, 50 % pulse, 100 % power, and continuous cooling by 0 °C water flow‐ through.

#### Preparation of libraries for the Illumina sequencing platform

Paired‐end libraries were prepared using the DNA sample kit (#1003382, Illumina) according to the manufacturer’s instructions, applying the following modifications: 200–220 bp fragments were cut out from an agarose gel after adapter ligation; right before library amplification uridine digestion was performed at 37 °C for 15’ in 5 μL 1x TE buffer, pH 7.5 with 1 U UNG (Applied Biosystems).

#### Sequencing

Amplified material was loaded onto channels of the flowcell at approximately 15 pM concentration. Sequencing was carried out using the 1G Illumina Genome Analyzer (Solexa) by running 51 paired‐end cycles according to the manufacturer’s instructions. Image deconvolution and quality value calculation was performed using the Goat module (Firecrest v.1.8.28 and Bustard v.1.8.28 programs) of the Illumina pipeline v.0.2.2.3.

### Read mapping

Read sequences of 51 bp were aligned to the zebra finch genome (tae.gut.3.2.4, August, 2008 [1]) using the read‐mapping algorithm TopHat version 1.0.8 with the following parameters: ‐p 4 (use 4 threads), ‐a 8min (anchor for mapping), ‐i 20 (minimum intron size), ‐I 20000 (maximum intron size), ‐mate‐std‐dev 20 (standard deviation of the mate distances), ‐mate‐inner‐dist 40 ‐F 0.05 (minimum fraction to call an isoform). Gene annotation was based on Ensembl v.55(ENS55). Further calculations were done using the statistical language R [10].

### Background noise estimation and expression

To avoid overestimation of background reads due to a lot of unknown transcribed units in inter‐ as well as intragenic regions, we generated a cumulative set of transcribed regions integrating all available external transcriptome information (ENS55 annotation, GeneBank mRNAs and ESTs, RefGene annotation, cross species alignments of cDNAs). As newly annotated regions we took all blocks larger than 1000 bp and at least supported by one cDNA evidence into account. 70 % of all mapped reads overlapped with this new annotation. Reads mapping outside of any mRNA/EST regions were considered as background, composed of intronic and intergenic regions.

Using the set of intronic and intergenic regions, we fitted a negative binomial model for experimental noise, which was used to assess expression of genes. Each ENS55 gene model was tested against this noise model and was declared as expressed if the number of overlapping reads was significantly (p<0.01) above background. The number of unique reads per gene was normalized by the total number of uniquely mapped reads per library and by the effective length of the gene (RPKM: reads per kbp per million) (Mortazavi et al., 2009). The coverage of individual exons was calculated using the following formula:

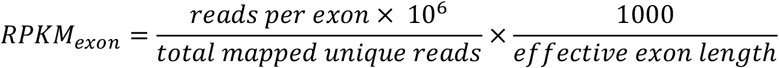

The effective length of an exon is the number of positions per exon, where reads can be aligned to without ambiguity (i.e. only unique positions).

### Annotation of new transcribed regions

Regions with two or more overlapping reads were called as blocks. Block‐coordinates were subsequently overlapped with the ENS55‐coordinates. If the region overlapped with the ENS55 annotation, it was kept unchanged for later processing. If it was a new region, we applied the following filter set to minimize false positives: length ≥ 80 bp, read count ≥ 3, coverage ≥ 2 RPKM. Regions that were not filtered out and were less than 50 bp apart from an existing ENS55 exon were merged with this ENS55 exon. An ENS55 exon was extended on the start or end coordinate if the extension was larger than 10 bp. After this initial round of calling transcribed regions, we integrated reads coming from splice junctions. For the ENS55‐supported blocks, the start and end coordinates were corrected according to the junction reads. The start‐ and end‐coordinates of the new blocks were changed if the junction reads ended not more than 10 bp within the block or 50 bp outside of the block. Even though we used a library preparation protocol retaining the directionality information [11], we did observe considerable signal on the opposite direction. We thus only kept new blocks with more than 50 % of the reads in the annotated direction.

### Analysis and statistics

Statistics and plots were generated with R and the ggplot2 package [10, 12]. The statistical test chosen for each data set is detailed in the results section of this thesis. Statistical properties of data shown in box plots are as follows: line in box (median), box (25^th^ and 75^th^ percentile), whisker (lowest or highest value within 1.5 * IQR (inter quartile range) of the 25^th^ or 75^th^ percentile), points below or above whiskers are outliers.

In density plots, the area under the curve corresponds to 1. The scale of the x‐axis depends on the bin‐width chosen to generate the plot. The default bin‐width is ‘range/30’. If two density plots were to be compared (**Figure 1C, Figure 3**), the same bin‐width was chosen for both plots.

**Figure 1:**
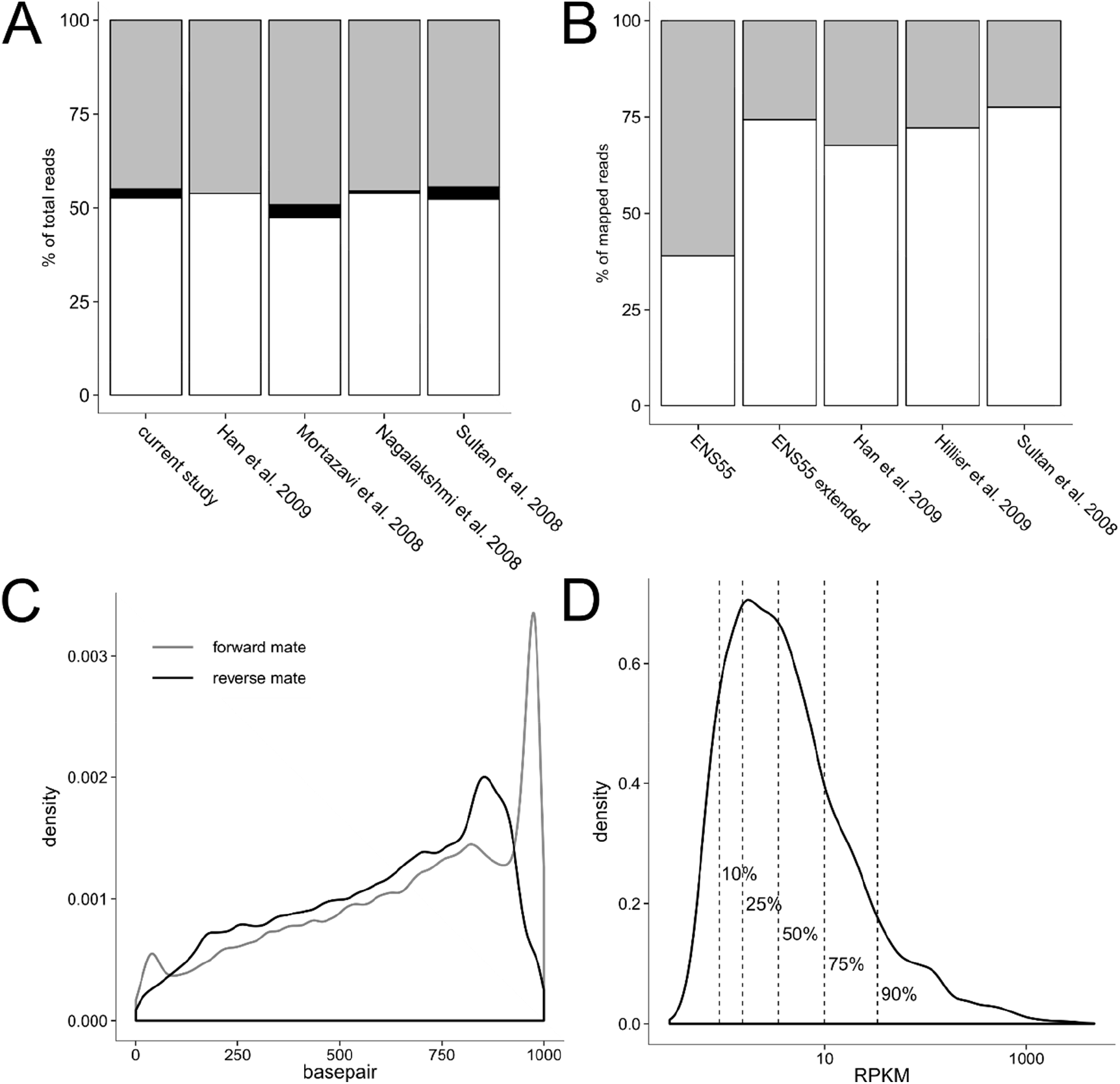
Mapping statistics. (A) Comparison of the read mapping success in different model organisms (from left to right: zebra finch, mouse, human, *S. cerevisiae*, human). In all species, more than 50 % of the reads mapped to the respective reference genome without gaps (white), a very small proportion of reads mapped to the genome in a gapped fashion, which indicates exon‐exon‐junctions (black). More than 40 % of all sequenced reads did not align to the reference genome (gray). (B) After overlapping the read‐coordinates with the annotation, the mapped reads were split into two groups: reads overlapping with the annotation (white) and reads in introns and intergenic regions (gray). After extending the zebra finch ENS55 annotation on the 3’‐end of gene models, the fraction of reads outside of the annotation was comparable to data from mouse, *C. elegans* and human. (C) Density plot of the relative position of reads on ENS55 transcripts, separated by mate of the paired end reads. (D) Distribution of the RPKM values of ENS55 gene models expressed above background (RPKM>0.4)(10%≙0.9RPKM, 25%≙1.5RPKM, 50%≙3.5RPKM, 75%≙9.9RPKM, 90%≙33.3RPKM).

## RESULTS

### Re‐annotation of the zebra finch genome

The standard workflow for high throughput investigation of gene expression typically requires a sequenced genome as well as an annotation of the transcribed regions. The first zebra finch genome, together with its annotation was published in 2010 [1]. An updated genome assembly was published by The Vertebrate Genome Project using new sequencing technologies on the DNA of the previously sequenced, male zebra finch [2]. The updated genome assembly included the W chromosome from a female sample (Blue55). Personal experience and communication with other researchers in the field indicates that, despite its improvement, the annotation is far from complete. Gene models are missing annotation of the UTRs, alternative splice forms are rarely annotated and genes might miss exons. We therefore decided to sequence the transcriptome of the zebra finch forebrain at two different ages and use the power of the data set to re‐annotate the zebra finch genome based on the ENS55 annotation.

#### Sequencing and mapping of reads

We carried out paired‐end sequencing of the polyA^+^‐RNA fraction from forebrains of male zebra finches at two behaviorally important time points: PHD50, in the middle of the sensorimotor period of song learning and PHD850, long after closure of the window for song learning. The mRNA of two pooled samples per stage was reverse transcribed into double stranded cDNA preserving the directionality of the mRNA and sequenced on the Illumina GAII platform [11]. Overall, 130 million 51 bp reads were generated from the 4 samples (30–35 million per sample) amounting to 6.6 Gigabases in total. Reads were aligned to the zebra finch genome (taeGut1) [1] using TopHat [13]. 52.55 % of all reads (68.5 million reads in total, 16–18 million reads per lane) mapped to the genome without gaps. At the time of analysis, the proportion of mapped reads was similar to the results in other, more established model organisms [14–17] (**Figure 1FiguA**). According to ENS55, 25.6 million bp of the genome are transcribed (2 % of the genome), thus all aligned reads together achieve an average depth of coverage of 34‐fold per transcribed base per sample.

### Gene expression analysis based on ENS55

Genes expressed in the male zebra finch forebrain were identified by overlapping the coordinates of ENS55 [18] with the coordinates of our mapped reads. Expression strength was calculated as reads per kilobase of transcript per million of sequenced reads (RPKM, [16]). Of the 68.5 million reads mapping to the genome, 38.95 % overlapped with the ENS55 annotation with at least one base pair. In comparison to results obtained with human, mouse or *C. elegans* data [14, 15, 19], the percentage of mapped reads within annotated regions was unusually low, indicating a large proportion of transcribed units were not annotated (**Figure 1B**). Inspection of the read distribution along the transcript revealed a bias in coverage towards the 3’‐end of transcripts (**Figure 1C**) as previously seen when sequencing oligo‐dT primed cDNA [14]. We consequently assumed that a large proportion of the reads mapping to unannotated regions in fact originated from the 3’‐UTR of transcripts, which was rarely annotated in the ENS55 zebra finch annotation. To test our hypothesis, we extended each ENS55 gene model by 3 kb on the 3’‐end. This simple extension increased the proportion of mapped reads within the annotation to levels (74.3 %, **Figure 1B**) comparable to studies on more established genetic model organisms, like mouse, *C. elegans* or human [14, 15, 19].

We used a negative binomial model to estimate background noise and tested each transcribed region (i.e. ENS gene) against it to decide if it was expressed (DABG p<0.01) or not (DABG p≤0.01). The average noise level was 1 read per 140 bp, which is within the range observed in other data sets [20]. Out of the 17,549 annotated genes, 10,475 were expressed in at least one of the four samples, with 9,869 and 10,103 genes for the juvenile and adult samples, respectively.

The majority of genes were expressed at low levels (55 % at RPKM<4), and conversely 0.63 % of the gens with the highest coverage accounted for 25 % of all reads overlapping the annotation. The level of expression spanned across four orders of magnitude in all samples, ranging from 0.6 to 4683 RPKM (**Figure 1D**).

### QPCR validation of gene expression quantification

In order to verify the expression strength for genes derived from the length‐normalized gene counts, we measured the expression of selected genes by QPCR. As template we used cDNA synthesized from the original enriched mRNA or total RNA, or total RNA extracted from the forebrains of six independent animals of the same ages as the sequenced animals. To select genes for the validation experiment, we divided the genes into three groups with either high (RPKM>32), medium (RPKM 4‐32) or low (RPKM<4) expression levels. Examples were selected from all three groups. We observed a good correlation between RNA‐seq and QPCR data on mRNA (7 genes, **Figure 2A**) as well as on the total RNA derived cDNA (7 genes, **Figure 2B**). As expected, the results were less correlated on the independent samples (**Figure 2C**).

**Figure 1:**
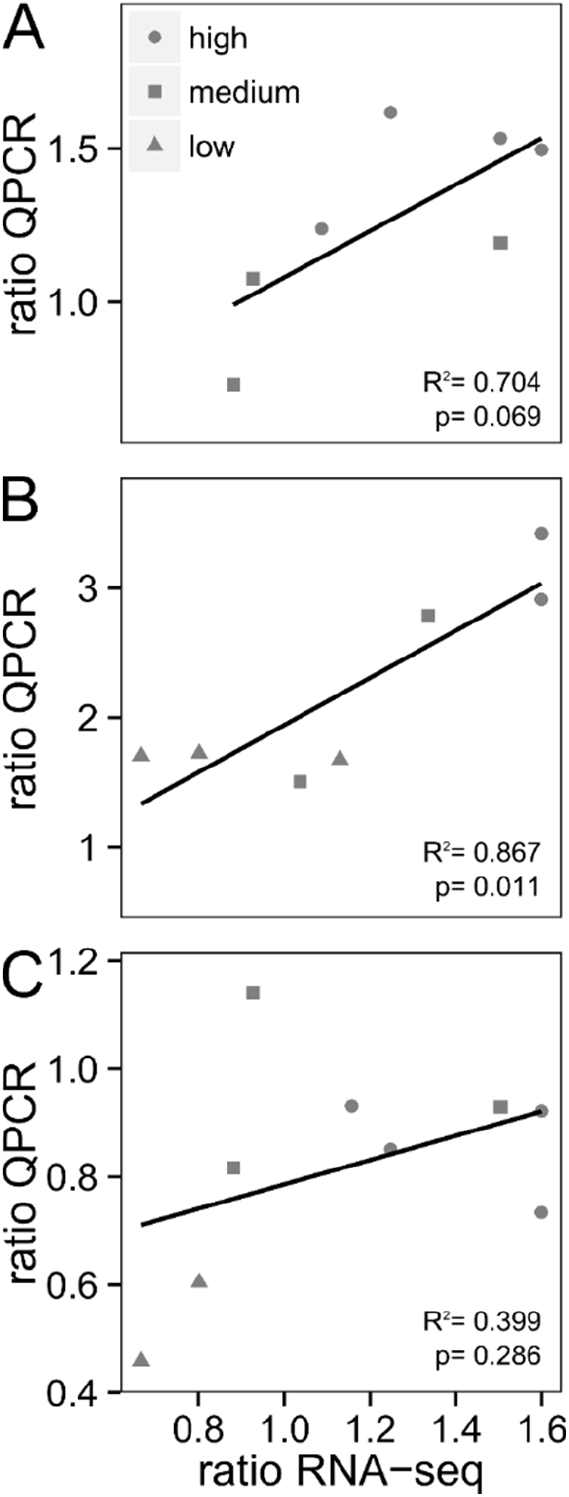
Comparison of gene quantification between RNA‐seq and QPCR. The ratio of expression between Old and Young was calculated for the RNA‐seq data as well as for the QPCR data. Results from RNA‐seq were compared with QPCR data from sequenced mRNA (A), sequenced total RNA (B), and total RNA from independent animals (n=3 per age, C). Each symbol refers to one gene. The shape of the symbol indicates the expression level of the gene according to RNA‐seq (high (circle), medium (square), low (triangle))

### Re‐annotation of transcribed regions

The ENS55 annotation of the zebra finch genome was primarily based on protein coding regions of the genome, thus 3’ and 5’‐UTRs were greatly underrepresented. Furthermore, the ENS 55 annotation reported 1.04 splice forms per gene on average, a much lower ratio than in other, well studied organisms. The lack of annotated splice forms is still prevalent. The current annotation (NCBI Taeniopygia guttata Annotation Release 106) lists an average of 2.28 splice forms per gene. For humans it is assumed, that almost all multi‐exon genes have at least one alternative splice form [20] and the most recent annotation (GCF_000001405.40‐RS_2023_10) of the human reference genome (GRCh38.p14). Our data provides high coverage information of the brain transcriptome with support for splice junctions coming from the reads aligning to two adjacent exons (junction reads). We thus decided to re‐annotate the transcribed regions of the zebra finch genome.

### Identifying transcribed regions

To identify transcribed regions, we started by searching for regions with at least two overlapping reads (henceforth called ‘blocks’), resulting in a total of 560,306 initial blocks for Old 1, 643,377 for Old 2, 578,911 for Young 1 and 643,377 for Young 2. Next, we divided the blocks into two groups: those overlapping with the ENS55 annotation and those not overlapping with the ENS55 annotation. The latter were subsequently filtered for a minimum RPKM of 2 and for at least three reads per block. Additionally, we set a minimum exon size of 80 bp, as less than 25 % of the ENS55 exons were smaller than 80 bp. We then pooled data from all sequenced lanes and merged all blocks less than 50 bp apart, because only 3.4 % of all ENS55 introns are smaller than 50 bp. In the next step the filtered blocks were merged with ENS55 regions if they were less than 50 bp apart. If a block elongated an existing ENS55 exon more than 10 bp, the coordinates were adjusted accordingly. These redefined ENS55‐blocks were subsequently also subjected to the abovementioned filters. To precisely annotate the boundaries of exons, all blocks were corrected using junction reads. Finally, we discarded all blocks with a read coverage mostly in the opposite direction than an annotated block.

### Exon re‐annotation

Our re‐annotation contained 154,690 exons, 89 % of which were supported by ENS55, while 11 % (16,922) were newly annotated. 28 % of the newly annotated regions were extensions of known ENS55 genes. Of the 137,768 ENS supported exons, 122,881 were unchanged in comparison to ENS55, while 14,887 exons were changed in length—all except one exon were extended. As expected, due to the 3’‐bias of our coverage and the underrepresented 3’‐UTR annotations in ENS55, we extended more ENS exons on the end than on the start (**Figure 3**). Additionally, the end extensions were longer than the start extensions (start: 1–5718 bp; end: 1– 12,370 bp). Most (70 %) of the end extended exons were annotated as the last exon of a gene. These exons had longer extensions than internal or first exons of a gene. This fact was very well demonstrated by the missing peak around 6 bp when only extensions of last exons are considered (**Figure 3**). The frequency distribution of start extensions is more homogeneous (**Figure 3**).

**Figure 3:**
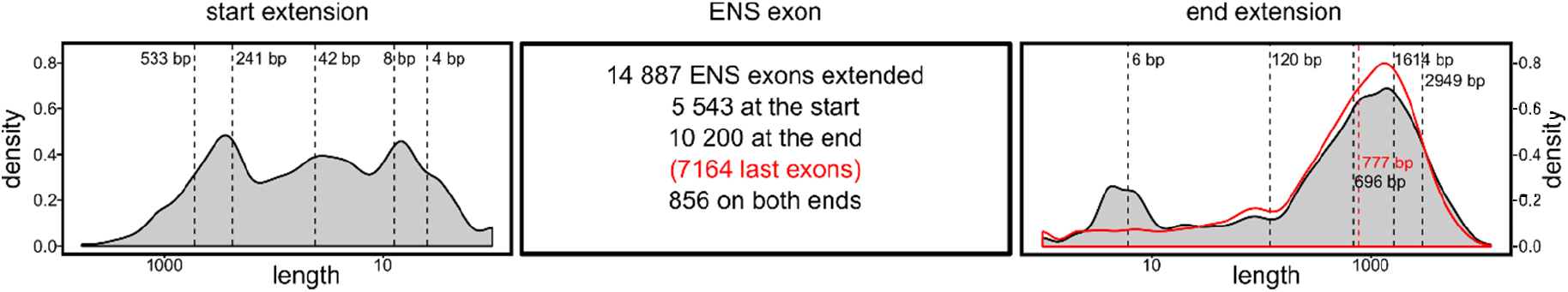
Extension of ENS exons. Depicted is the distribution of start and end extensions of ENS55 exons over their length. Extensions at the end of exons occurred more often and were longer than start extensions, which was mostly due to the annotation of putative 3’‐UTRs for the last exons of a gene model (red density plot). Dashed lines represent the 10%, 25%, 50%, 75% and 90% quantiles (left to right for end‐extensions, right to left for start extensions). The red density plot and red dashed line represent the end extensions of last exons and the median, respectively. Binwidth = 22 for all plots.

### New gene models

Our annotation contained 25,176 gene models. 11,391 of those were new. Of the 13,875 gene models supported by the ENS55 annotation, 2,822 were extended by new exons. The gene models unsupported by ENS55 consisted of one up to eleven exons, with the vast majority (10,966) containing only one exon. The ENS55 supported gene models were mostly unchanged compared to the original ENS55 annotation. In 419 cases we concatenated two to seven ENS55 genes into one gene model.

### PCR validation of re‐annotation events

Additional confirmation of the results of our re‐annotation procedure was achieved by PCR based verification of re‐annotation events on six independent samples (**Figure 4‐8**). In most cases the re‐annotation and PCR verification were in accordance. Deviations between the re‐annotation and manual curation are attributed to factors like filtering steps in the re‐annotation pipeline or gaps in the genome assembly.

**Figure 4:**
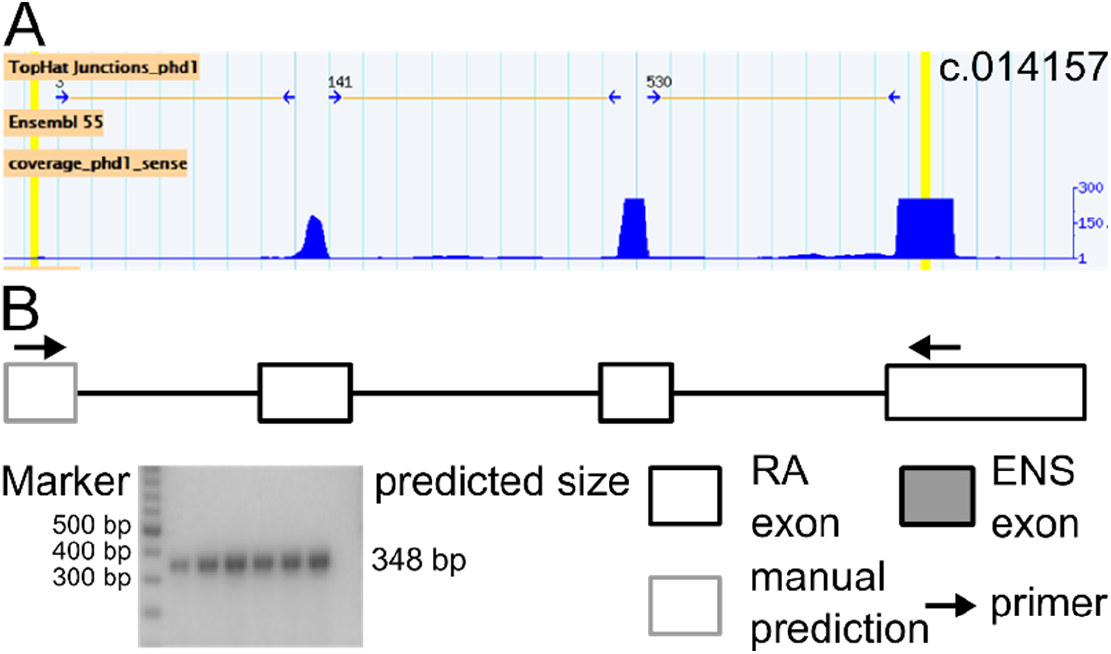
PCR validation of component c.014157. (A) Genome browser representation of the read coverage (blue track) and junction reads (horizontal yellow lines). Bright yellow vertical bars mark the primer binding sites for the PCR verification. (B) Exon‐intron structure of the newly annotated component c.014157. Boxes represent exons, lines the possible exon‐exon‐connections. Arrows indicate the position of the primers used for PCR‐validation. The fill of the box indicates if the exon was annotated in ENS55 (gray). The color of the outline indicates if the exon was annotated by the re‐annotation pipeline (black) or manually predicted (gray) from the data. The agarose gel shows the size of the amplified product. cDNA template originated from the forebrain of young (first three lanes) and old (next three lanes) animals. The size marker was run on the first lane and is annotated on the left, the predicted fragment size on the right. The last lane is the product from the NTC.

### Newly annotated components

We chose two entirely new gene models, c.014157 (**Figure 4**) and c.008232 (**Figure 5**), consisting of more than one exon for PCR verification. In the case of c.014157, three exons were annotated and connected into a new component. Manual inspection of the locus provided evidence for a fourth exon connected to the 5’‐end of the component, which could be confirmed by PCR (**Figure 4**). Component c.008232 was annotated with four exons lacking alternative splicing events. Manual curation of the annotation revealed evidence for three additional upstream exons, which were alternatively spliced into three possible transcripts. PCR verification confirmed the re‐annotation as well as the manual curation of the component (**Figure 5**). The three exons that were not annotated by our re‐annotation due to our filtering steps for coverage above 2 RPKM, a minimum exon size of 80 bp and overlap with a gap in the genome assembly.

**Figure 5:**
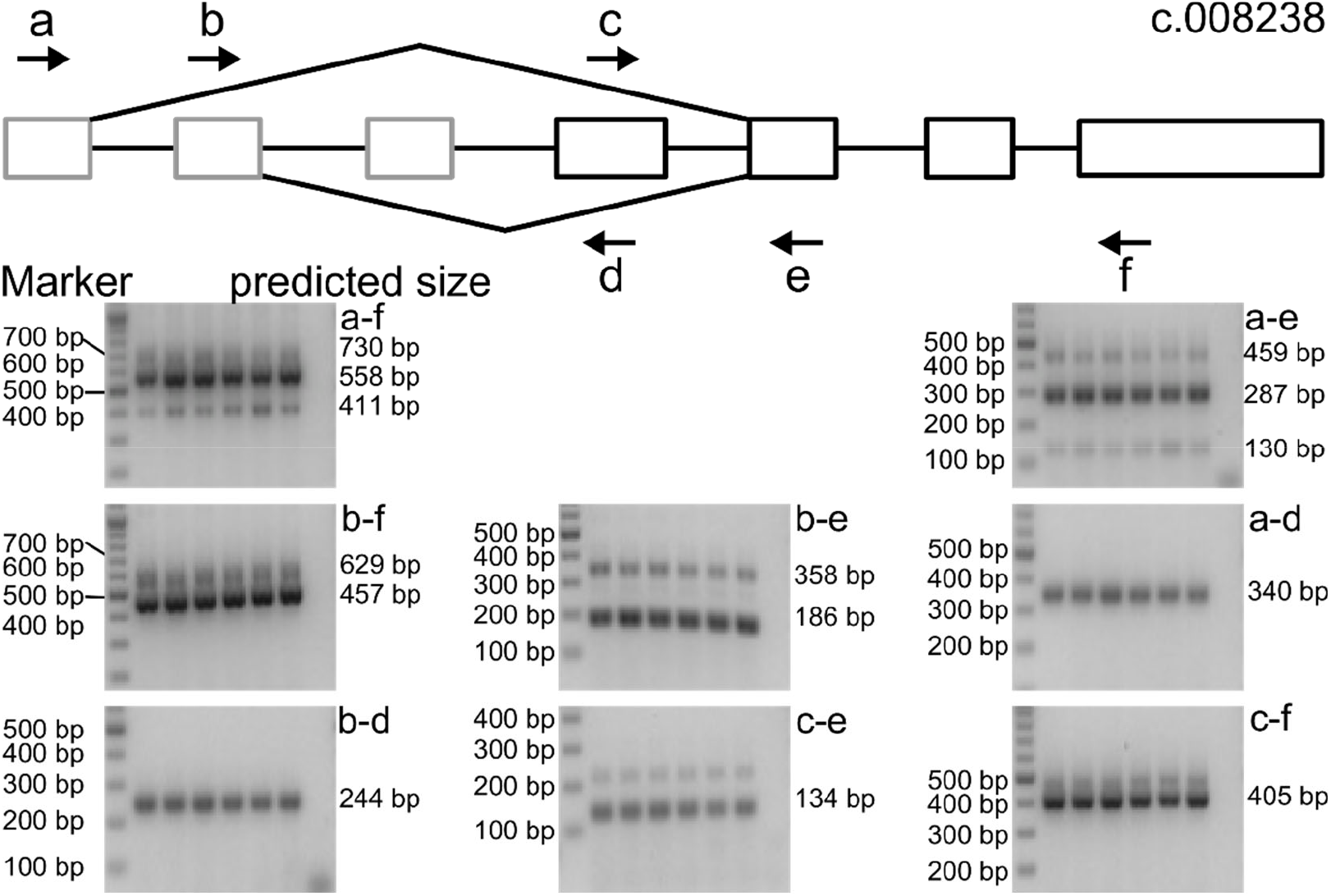
PCR validation of component c.008232. The component was annotated with four exons. Three additional exons were manually detected and connected to the component. PCR validation confirmed all predictions. When using primer c, a second amplicon, longer than predicted, was amplified. The transcriptome data did not provide any explanation for this. Mispriming could not be excluded in this case. The agarose gel shows the size of the amplified product. cDNA template originated from the forebrain of young (first three lanes) and old (next three lanes) animals. The size marker was run on the first lane and is annotated on the left, the predicted fragment size on the right. The last lane is the product from the NTC.

### 3’‐UTR extensions

The zebra finch ENS55 annotation rarely annotated UTR‐regions of gene models. Due to a coverage bias towards the 3’‐end of transcripts (**Figure 1C**), we were able to re‐annotate more than 7,000 last exons by extending the end‐coordinate. Two of these extensions were confirmed by PCR (**Figure 6, Figure 7 PCR e–i**). In both cases a fragment of the predicted length was amplified. In the case of *OLFM1* (**Figure 7**) the extension could only be confirmed for samples originating from old individuals. The transcriptome data did not indicate an age‐dependent difference in UTR length for *OLFM1*. As the signal strength for the long UTR extension PCRs was always weaker than that for shorter PCRs, this finding could be an effect of amplification efficiencies of long fragments.

**Figure 6:**
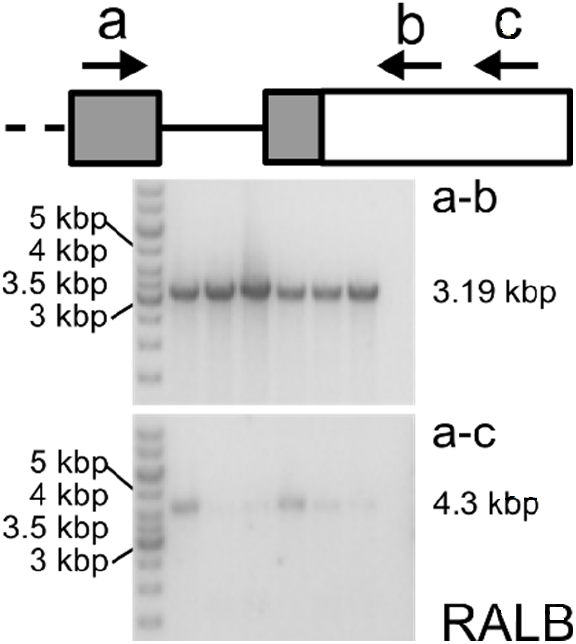
PCR validation of RALB 3’‐UTR extension. The last ENS exon of the ENS55 annotation was extended by more than 3 kbp, which was supported by PCR. The agarose gel shows the size of the amplified product. cDNA template originated from the forebrain of young (first three lanes) and old (next three lanes) animals. The size marker was run on the first lane and is annotated on the left, the predicted fragment size on the right. The last lane is the product from the NTC.

**Figure 7:**
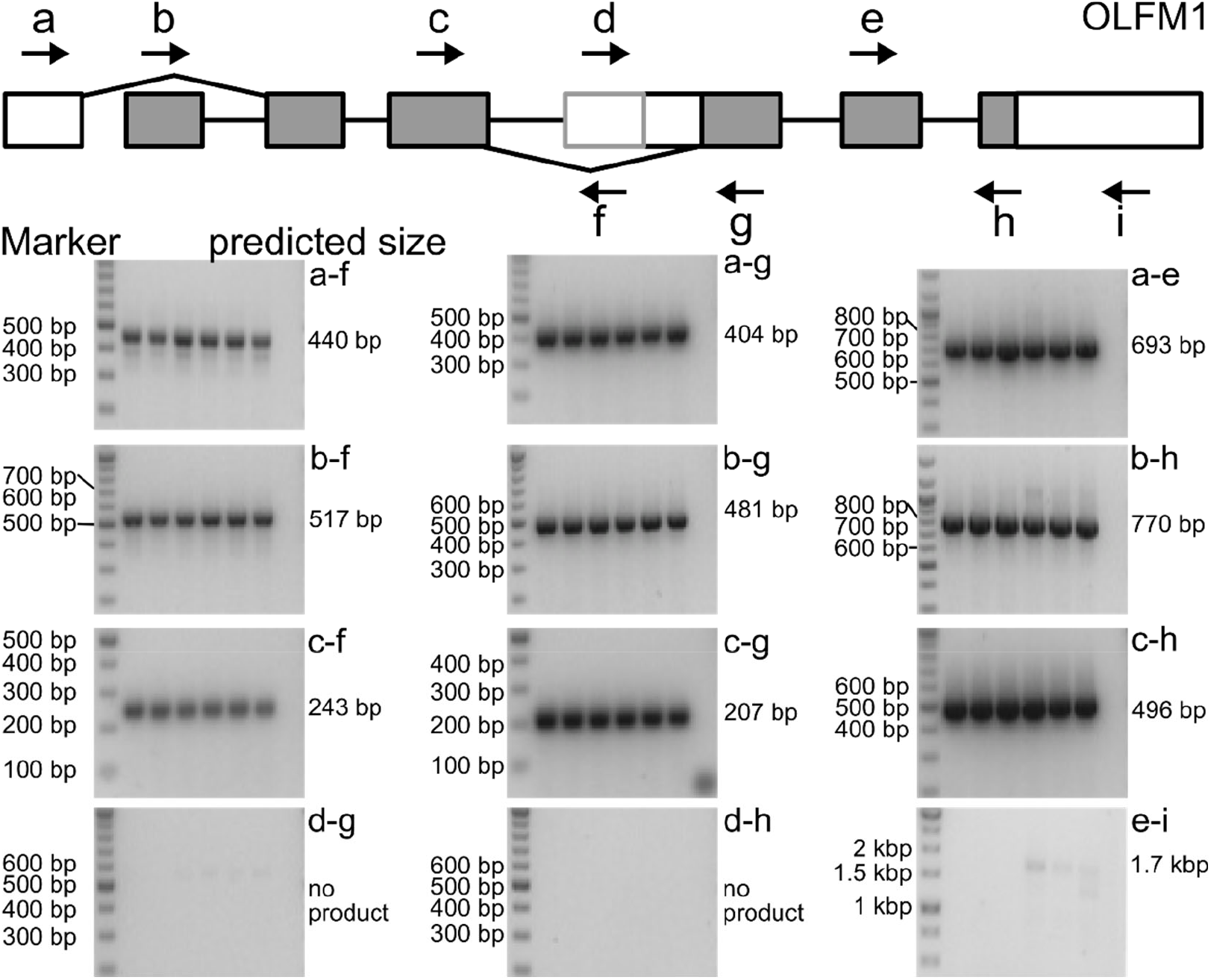
PCR validation of *OLFM1* re‐annotation. The *OLFM1* gene was re‐annotated with one new first exon, a large exon extension of an internal exon as well as a 3’‐UTR extension of the last exon. The new first exon and 3’‐UTR extension could be confirmed by PCR. However, manual curation of the internal exon extension revealed that it is in fact a new alternative last exon, which was fused to the start of an internal ENS exon. The fusion was not supported by PCR. cDNA template originated from the forebrain of young (first three lanes) and old (next three lanes) animals. The size marker was run on the first lane and is annotated on the left, the predicted fragment size on the right. The last lane is the product from the NTC.

### New exons

Our re‐annotation data set contained 4,797 new exons, which extended the already known ENS55 genes. One example was the extension of the *OLFM1* gene by an alternative start exon (**Figure 7**). The gene coding for *Olfm1* in mice (*mus musculus*) contains two alternative first and last exons, resulting in 4 possible transcripts. The re‐annotation data set did not contain an alternative last exon for *OLFM1*, but extended the start coordinates of one internal exon by several hundred base pairs (**Figure 7**, internal long exon depicted by 3 boxes). Closer inspection of the raw read data revealed that the new alternative last exon was fused with the downstream ENS exon, resulting in the start coordinate extension of the aforementioned ENS exon. However, the read distribution did not support fusing the new last exon and the known ENS exon, as no reads spanned over both and no junction reads were aligned to this junction. PCR results supported the manual curation.

### Alternative splicing

Alternative splicing events, leading to the generation of more than one possible transcript per gene are rarely annotated by ENS55 for zebra finches, which is reflected in the low number of transcripts per gene (1.04). Manual screening of the transcriptome data revealed support for a multitude of unannotated alternative splicing events, many of which are known from mouse and human data. One example is a skipped exon in the *KAT2B* gene. Primers were designed to bind to the exons up‐ and downstream of the skipped exon. PCR amplification confirmed the presence of two amplicons of the predicted size (**Figure 8**). An alternative example is the *OLFM1* gene, which can be spliced into four distinct transcripts by using two alternative first and last exons (**Figure 7**). All four transcripts were detected by PCR.

**Figure 8:**
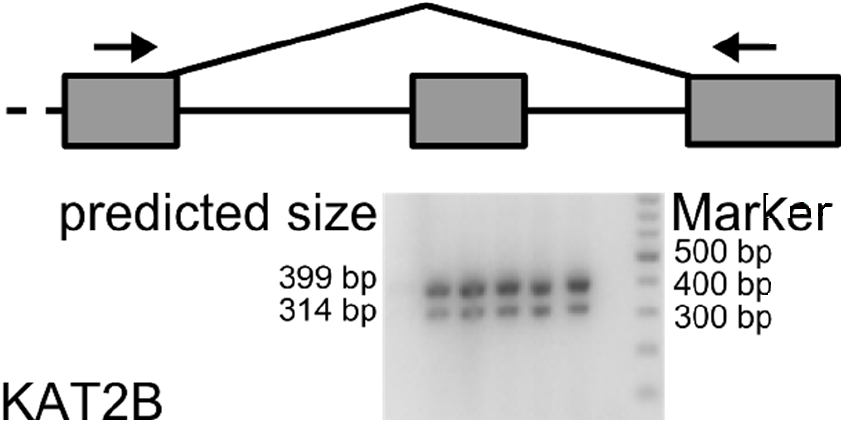
*KAT2B* exon skipping. PCR confirmed the skipping of an internal exon of the KAT2B gene. The alternative splicing event is described for the mouse homologue. cDNA template originated from the forebrain of young (first three lanes) and old (next three lanes) animals. The size marker was run on the first lane and is annotated on the left, the predicted fragment size on the right. The last lane is the product from the NTC.

### Quantification of gene expression using the new annotation

Changes in genome annotation could have the potential to affect gene quantification. In most RNA‐seq experiments read distribution over the length of transcripts is not uniform. This bias depends in large parts on the priming strategy in the cDNA preparation step. For example, using oligo‐dT primers introduces a 3’‐bias. Thus, the length of the annotated 3’‐UTR can have a large influence on the expression strength calculated for a given gene, which is of special importance when comparing the expression levels across genes in a data set. To test if the quantification of gene expression changed due to the re‐annotation of gene models, we compared the RPKM values derived from the ENS55 and new annotation with each other (**Figure 9**). The Pearson‐ correlation coefficient between both data sets was 0.89 for the ‘Old’ samples. The shape of the distribution indicates that the RPKM values derived from the new annotation tended to be higher compared with the RPKM values derived from the ENS55 annotation, which is reflected in a higher mean, median and maximum value of the new quantification.

**Figure 9:**
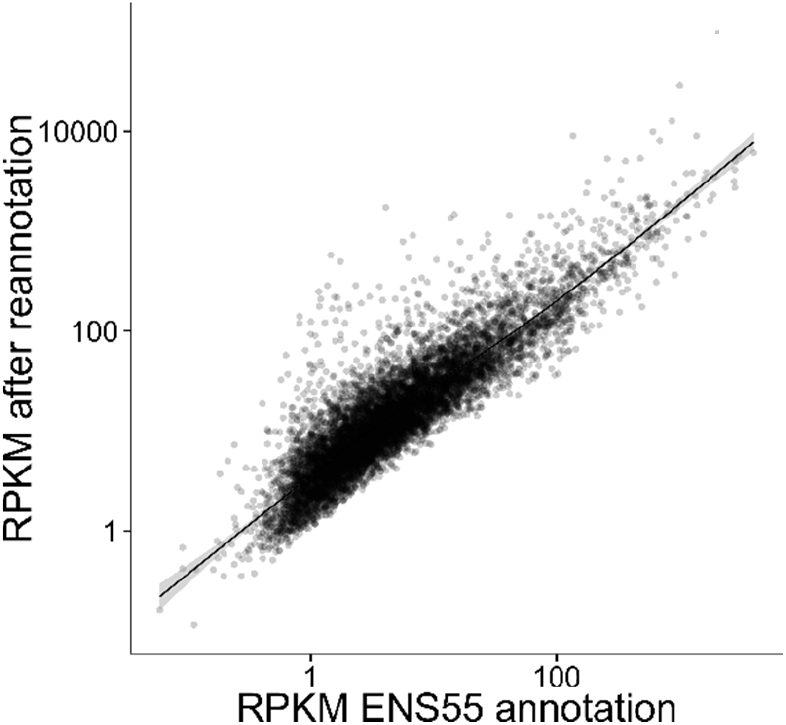
Comparison of gene expression quantification. Mean RPKM values of Old 1 and Old 2 were calculated using the ENS55 annotation or the new annotation and plotted against each other. Only components encompassing a single ENS55 gene are depicted. Both axes are plotted in log‐scale to accommodate the range of the data. Each dot represents one gene, the gray area around the fitted line denotes the 0.95 confidence interval.

## DISCUSSION

We successfully used transcriptome data derived from of the male zebra finch forebrain to annotate new gene models, extend known gene models with new exons and extend known exons, especially last exons, by annotating putative 3’‐UTRs. We further confirmed some of these re‐annotation events experimentally to validate our results.

The experimental validation of our re‐annotation efforts generally resulted in good agreement between the predicted data and the PCR results. However, manual curation of the loci usually improved the prediction and could always be confirmed by PCR. The three reasons for discrepancies between the re‐annotation and the PCR verification were (1) genome quality, (2) high coverage on intronic regions and (3) filtering artefacts of the annotation pipeline. The quality of the reference genome can greatly affect the success of transcriptome sequencing projects in the mapping stage because signal coming from incorrectly assembled or missing regions is lost in genome guided transcript reconstruction even if a transcribed region was sequenced. Two examples for such cases are the VLDLR gene [5] and the newly annotated gene model c.008238, both of which were missing an exon or the correct exon boundary due to gaps in the genome build. Another factor confounding the annotation of transcripts is the number of reads falling into intronic regions. We defined a transcribed block as a continuous stretch of the genome with read coverage above 2 RPKM. In the initial block calling step, we did not impose any filter for coverage uniformity along the length of the block, thus introns with read coverage above 2 RPKM were merged with the surrounding exons. The occurrence of reads in intronic regions is well described and especially prominent in the brain [21], with more than 20 % of the reads mapping to intronic regions in the human brain. These non‐exonic reads originate from the so‐called ‘dark matter’ RNA, which by definition is RNA with no understood function [21, 22] and often originates from introns [23]. Because they can be reproducibly detected, these intronic RNAs are considered to be functionally relevant. Their function probably lies in the perturbation of the effect of regulatory non‐coding RNAs. Even though we implemented a step to filter out exons with less than 10 % coverage of the main exon we still annotate intronic regions in some cases. Whether these are artefacts or of biological importance cannot be inferred from our data.

In summary, we added a substantial number of new genes, exons and exon extensions to the ENS55 zebra finch annotation, which illustrates the need and value to incorporate new transcriptional evidence into existing annotations.

## ACKNOWLEDGMENTS

We would like to thank Ursula Kobalz for technical support and Janett Meinecke for animal care.

## DATA AND CODE AVAILABILITY

Data and code is available from the corresponding author on reasonable request.

## Notes

### Competing Interest Statement

The authors have declared no competing interest.

## REFERENCES

1. Warren WC, Clayton DF, Ellegren H, Arnold AP, Hillier LW, Kunstner A, et al. The genome of a songbird. Nature. 2010;464(7289):757–62. Epub 2010/04/03. 10.1038/nature08819. PubMed PMID: 20360741; PubMed Central PMCID: PMCPMC3187626.

2. Korlach J, Gedman G, Kingan SB, Chin CS, Howard JT, Audet JN, et al. De novo PacBio long-read and phased avian genome assemblies correct and add to reference genes generated with intermediate and short reads. Gigascience. 2017;6(10):1–16. Epub 2017/10/13. 10.1093/gigascience/gix085. PubMed PMID: 29020750; PubMed Central PMCID: PMCPMC5632298.

3. Kosubek-Langer J, Scharff C. Dynamic FoxP2 levels in male zebra finches are linked to morphology of adult-born Area X medium spiny neurons. Sci Rep. 2020;10(1):4787. Epub 2020/03/18. 10.1038/s41598-020-61740-6. PubMed PMID: 32179863; PubMed Central PMCID: PMCPMC7075913.

4. Norton P, Barschke P, Scharff C, Mendoza E. Differential song deficits after lentivirus-mediated knockdown of FoxP1, FoxP2 or FoxP4 in Area X of juvenile zebra finches. J Neurosci. 2019. Epub 2019/10/24. 10.1523/JNEUROSCI.1250-19.2019. PubMed PMID: 31641053.

5. Adam I, Mendoza E, Kobalz U, Wohlgemuth S, Scharff C. FoxP2 directly regulates the reelin receptor VLDLR developmentally and by singing. Mol Cell Neurosci. 2016;74:96–105. Epub 2016/04/24. 10.1016/j.mcn.2016.04.002. PubMed PMID: 27105823.

6. Zhao W, Garcia-Oscos F, Dinh D, Roberts TF. Inception of memories that guide vocal learning in the songbird. Science. 2019;366(6461):83–9. Epub 2019/10/12. 10.1126/science.aaw4226. PubMed PMID: 31604306.

7. Medina CA, Vargas E, Munger SJ, Miller JE. Vocal changes in a zebra finch model of Parkinson’s disease characterized by alpha-synuclein overexpression in the song-dedicated anterior forebrain pathway. PLoS One. 2022;17(5):e0265604. Epub 2022/05/05. 10.1371/journal.pone.0265604. PubMed PMID: 35507553; PubMed Central PMCID: PMCPMC9067653.

8. Liu WC, Kohn J, Szwed SK, Pariser E, Sepe S, Haripal B, et al. Human mutant huntingtin disrupts vocal learning in transgenic songbirds. Nat Neurosci. 2015;18(11):1617–22. Epub 2015/10/06. 10.1038/nn.4133. PubMed PMID: 26436900.

9. Agate RJ, Scott BB, Haripal B, Lois C, Nottebohm F. Transgenic songbirds offer an opportunity to develop a genetic model for vocal learning. Proc Natl Acad Sci U S A. 2009;106(42):17963–7. Epub 2009/10/10. 10.1073/pnas.0909139106. PubMed PMID: 19815496; PubMed Central PMCID: PMCPMC2764872.

10. R Core Team. R: A Language and Environment for Statistical Computing. R Foundation for Statistical Computing, Vienna, Austria. 2013.

11. Parkhomchuk D, Borodina T, Amstislavskiy V, Banaru M, Hallen L, Krobitsch S, et al. Transcriptome analysis by strand-specific sequencing of complementary DNA. Nucleic Acids Res. 2009. Epub 2009/07/22. doi: gkp596 [pii] 10.1093/nar/gkp596. PubMed PMID: 19620212.

12. Wickham H. ggplot2: elegant graphics for data analysis: Springer Publishing Company, Incorporated; 2009 2009.

13. Trapnell C, Pachter L, Salzberg SL. TopHat: discovering splice junctions with RNA-Seq. Bioinformatics. 2009. Epub 2009/03/18. doi: btp120 [pii] 10.1093/bioinformatics/btp120. PubMed PMID: 19289445.

14. Han X, Wu X, Chung WY, Li T, Nekrutenko A, Altman NS, et al. Transcriptome of embryonic and neonatal mouse cortex by high-throughput RNA sequencing. Proc Natl Acad Sci U S A. 2009;106(31):12741–6. Epub 2009/07/21. doi: 0902417106 [pii] 10.1073/pnas.0902417106. PubMed PMID: 19617558.

15. Sultan M, Schulz MH, Richard H, Magen A, Klingenhoff A, Scherf M, et al. A global view of gene activity and alternative splicing by deep sequencing of the human transcriptome. Science. 2008;321(5891):956–60. Epub 2008/07/05. doi: 1160342 [pii] 10.1126/science.1160342. PubMed PMID: 18599741.

16. Mortazavi A, Williams BA, McCue K, Schaeffer L, Wold B. Mapping and quantifying mammalian transcriptomes by RNA-Seq. Nat Methods. 2008;5(7):621–8. Epub 2008/06/03. doi: nmeth.1226 [pii] 10.1038/nmeth.1226. PubMed PMID: 18516045.

17. Nagalakshmi U, Wang Z, Waern K, Shou C, Raha D, Gerstein M, et al. The transcriptional landscape of the yeast genome defined by RNA sequencing. Science. 2008;320(5881):1344–9. Epub 2008/05/03. doi: 1158441 [pii] 10.1126/science.1158441. PubMed PMID: 18451266.

18. Flicek P, Aken BL, Ballester B, Beal K, Bragin E, Brent S, et al. Ensembl’s 10th year. Nucleic Acids Res. 2010;38(Database issue):D557–62. 10.1093/nar/gkp972. PubMed PMID: 19906699; PubMed Central PMCID: PMC2808936.

19. Hillier LW, Reinke V, Green P, Hirst M, Marra MA, Waterston RH. Massively parallel sequencing of the polyadenylated transcriptome of C. elegans. Genome Res. 2009;19(4):657–66. Epub 2009/02/03. doi: gr.088112.108 [pii] 10.1101/gr.088112.108. PubMed PMID: 19181841.

20. Wang ET, Sandberg R, Luo S, Khrebtukova I, Zhang L, Mayr C, et al. Alternative isoform regulation in human tissue transcriptomes. Nature. 2008;456(7221):470–6. Epub 2008/11/04. doi: nature07509 [pii] 10.1038/nature07509. PubMed PMID: 18978772; PubMed Central PMCID: PMC2593745.

21. Kapranov P, St Laurent G, Raz T, Ozsolak F, Reynolds CP, Sorensen PH, et al. The majority of total nuclear-encoded non-ribosomal RNA in a human cell is ‘dark matter’ un-annotated RNA. BMC Biol. 2010;8:149. 10.1186/1741-7007-8-149. PubMed PMID: 21176148; PubMed Central PMCID: PMC3022773.

22. van Bakel H, Nislow C, Blencowe BJ, Hughes TR. Most “dark matter” transcripts are associated with known genes. PLoS Biol. 2010;8(5):e1000371. 10.1371/journal.pbio.1000371. PubMed PMID: 20502517; PubMed Central PMCID: PMC2872640.

23. St Laurent G, Shtokalo D, Tackett MR, Yang Z, Eremina T, Wahlestedt C, et al. Intronic RNAs constitute the major fraction of the non-coding RNA in mammalian cells. BMC Genomics. 2012;13:504. 10.1186/1471-2164-13-504. PubMed PMID: 23006825; PubMed Central PMCID: PMC3507791.

